# hiPSC-CM Electrophysiology: Impact of Temporal Changes and Study Parameters on Experimental Reproducibility

**DOI:** 10.1101/2023.10.02.560475

**Authors:** Devon Guerrelli, Jenna Pressman, Nikki Posnack

## Abstract

Human induced pluripotent stem cell-derived cardiomyocytes (hiPSC-CMs) are frequently used for preclinical cardiotoxicity testing and remain an important tool for confirming model-based predictions of drug effects in accordance with the Comprehensive *in Vitro* Proarrhythmia Assay (CiPA) initiative. Despite the considerable benefits hiPSC-CMs provide, concerns surrounding experimental reproducibility have emerged. Our study aimed to investigate the effects of temporal changes and experimental parameters on hiPSC-CM electrophysiology. hiPSC-CMs (iCell cardiomyocyte^2^) were cultured for 14 days and biosignals were acquired using a microelectrode array (MEA) system. Continuous recordings revealed a 22.6% increase in the beating rate and 7.7% decrease in the field potential duration (FPD) during a 20-minute equilibration period. Location specific differences across a multiwell plate were also observed, with hiPSC-CMs in the outer rows beating 8.8 beats per minute (BPM) faster than the inner rows. Cardiac endpoints were also impacted by cell culture duration; from 2-14 days the beating rate decreased (–12.7 BPM), FPD lengthened (+257 ms), and spike amplitude increased (+3.3 mV). Cell culture duration (4-10 days) also impacted hiPSC-CM drug responsiveness (E-4031, nifedipine, isoproterenol). Our study highlights multiple sources of variability that should be considered and addressed when performing hiPSC-CM MEA studies. To improve reproducibility and data interpretation, MEA-based studies should establish a standardized protocol and report key experimental conditions (e.g., culture time, equilibration time, electrical stimulation settings, report raw data values).

**New & Noteworthy:** We demonstrate that hiPSC-CM electrophysiology measurements are significantly impacted by slight deviations in experimental techniques including electrical stimulation protocols, equilibration time, well-to-well variability, and length of hiPSC-CM culture. Furthermore, our results indicate that hiPSC-CM drug responsiveness changes within the first two weeks following defrost.

Graphical Abstract Image created with Biorender.com

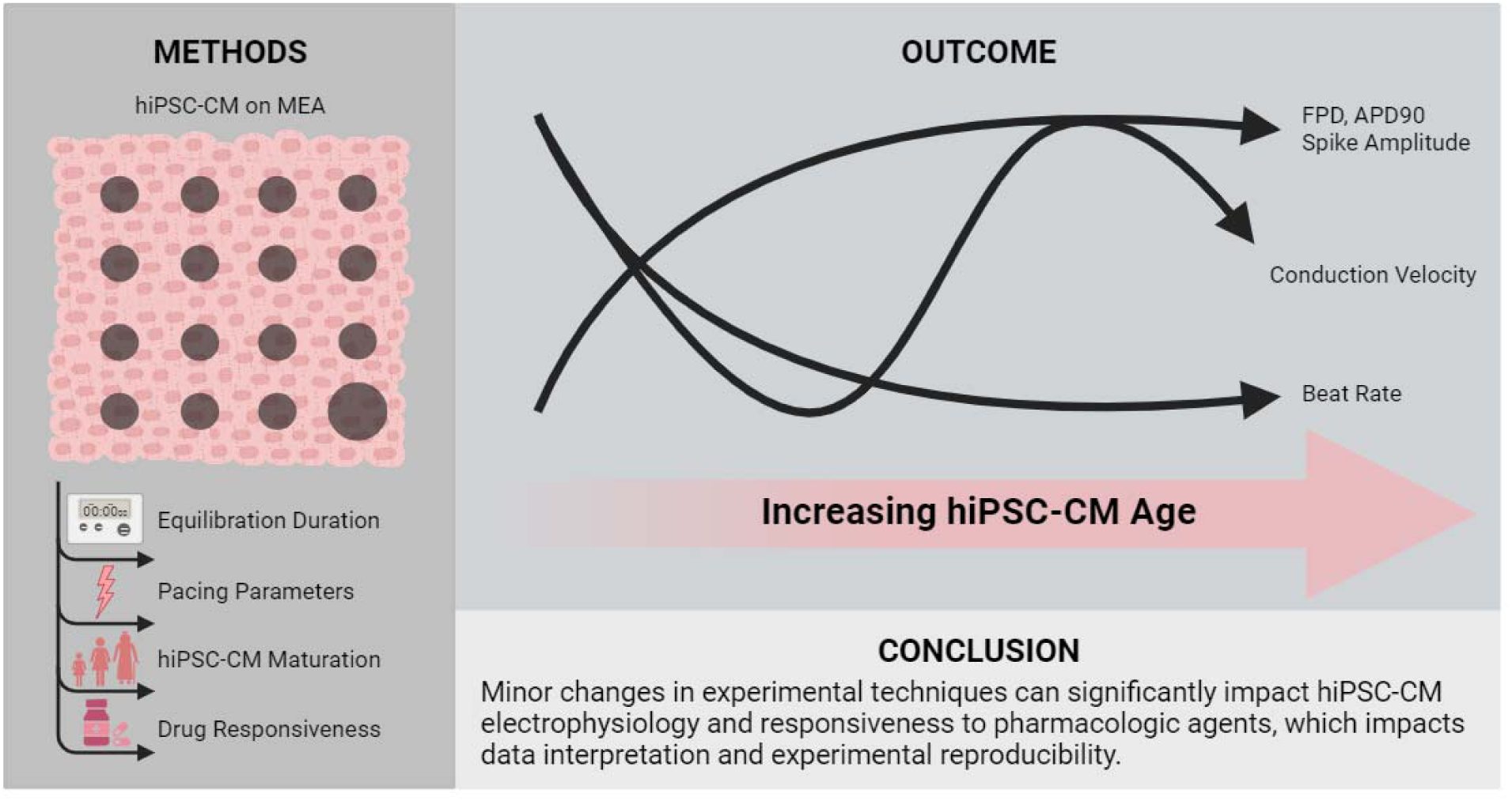

## Introduction

Human induced pluripotent stem cell-derived cardiomyocytes (hiPSC-CMs) have been widely adopted as a tool for translational research studies focused on the efficacy, safety, and cardiotoxicity of drug compounds and environmental chemicals. As previously reviewed (1–4), hiPSC-CMs offer several advantages to the cardiac research field: 1) hiPSC-CM are a renewable source of human-derived tissue, 2) hiPSC-CM are sensitive and responsive to drugs/chemicals that cause cardiotoxicity through structural, electrophysiology, or contractile disturbances, 3) patient-derived hiPSC-CM can be generated for disease modeling and precision medicine, and 4) hiPSC-CM more accurately recapitulate human cardiac biology, as compared to other animal models. While animal models may fail to predict drug safety or efficacy due to species-specific differences (5, 6) – hiPSC-CM express similar cardiac ion channels as human cardiomyocytes, have an action potential duration that is comparable to the human electrocardiographic QT interval, and hiPSC-CM are highly responsive to ion channel blocking drugs (7–13). One limitation to using an hiPSC-CM model is the immature phenotype of these cells (e.g., small size, spontaneous beating activity (14, 15)); although, multiple technical strategies have been identified to effectively mature these cells to more closely mimic adult human cardiomyocytes (16–21).

Given these advantages, hiPSC-CM have been incorporated as an integral tool in the Comprehensive in Vitro Proarrhythmia (CiPA) initiative, which was established to more precisely identify drug compounds with proarrhythmic risk using nonclinical assays (22, 23). Indeed, hiPSC-CMs were shown to accurately predict the proarrhythmic potential of ∼85% of drugs (26 tested) known to cause clinical QT prolongation (24). Further, hiPSC-CMs were both sensitive (87%) and specific (70%) when used to screen 51 drug compounds for their effects on cardiac contraction (25). Such large-scale cardiotoxicity screening assays can be performed using a high-throughput microelectrode array (MEA) platform, which allows one to measure both electrical and contractile properties of hiPSC-CM. Briefly, hiPSC-CMs are plated atop microelectrodes embedded within a cell culture plate, which allows for continuous (non-invasive) recordings of: 1) extracellular field potentials akin to an electrocardiogram, 2) local extracellular action potentials akin to the cardiac action potential, and 3) impedance signals corresponding to cardiomyocyte contractility (26–28). Although hiPSC-CM MEA studies can be used to identify cardiotoxic compounds, multiple studies have reported some inconsistences in the acquired data across study sites and cell lines (24, 29–35). Multiple factors are likely to contribute to such data discrepancies, including phenotypic differences in each individual hiPSC-CM line used, cell culture conditions (e.g., plating density, duration of cell culture, serum vs serum-free media), and the recording protocol implemented (e.g., gas/temperature control, length of equilibration time, pacing parameters). For example, the hiPSC-CM spontaneous beating rate is influenced by both the recording temperature and plating density (36–38), and cells with a slower beating rate can have an exaggerated pro-arrhythmic response to Ikr blockers (31). With this mind, we aimed to investigate experimental factors that may influence hiPSC-CM MEA study results. Controlling for and clearly defining study parameters that can influence MEA results will help to improve data interpretation and experimental reproducibility between labs, which is an increasing focus of the cardiovascular research field (39).

## Methods

### Culture and maintenance of hiPSC-CMs

MEA electrodes (#M384-tMEA-24W & #M384-BIO-24; Axion Biosystems) were coated with 50 µg/mL fibronectin (#33010018; Gibco) diluted in sterile phosphate buffered saline (#10010023; Gibco) for 1 hour in a cell culture incubator (37°C, 5% CO_2_). Subsequently, fibronectin was removed and iCell cardiomyocytes^2^ (#01434; Fujifilm Cellular Dynamics) were plated atop a 16-electrode array at a density of 40–65,000 cells per well in iCell cardiomyocyte plating media (#M1001; Fujifilm Cellular Dynamics). hiPSC-CMs were maintained in a cell culture incubator for 1-3 hours to allow cells to adhere to the electrode array, and thereafter, the media was removed and replaced with iCell serum-based maintenance media (#M1003; Fujifilm Cellular Dynamics) supplemented with 2 µg/mL ciprofloxacin (#17850-5G-F; Sigma-Aldrich). A full volume media change occurred 1 day post defrost; half volume media changes occurred every 2-3 days for the remainder of the experimental period.

### Microelectrode array recordings

*Equilibration:* A Maestro Edge MEA and impedance system (Axion BioSystems) was used for all hiPSC-CM physiological measurements; all experiments were conducted at 37°C with 5% CO_2_. To assess phenotypic changes that may occur during equilibration to the MEA system, cellular recordings were acquired continuously for 20 minutes while hiPSC-CMs were beating spontaneously. For all subsequent experiments, hiPSC-CMs were allowed to equilibrate to the MEA system for 20 minutes to reach stead state conditions. Thereafter, cellular recordings were acquired both while hiPSC-CMs were beating spontaneously and also in response to external pacing.

*MEA parameters of interest*: Biosignals were acquired from hiPSC-CMs plated atop a 16-electrode array (**Figure 1**). Electrophysiology parameters of interest included the spontaneous beating rate, field potential duration (FPD), spike amplitude, and conduction velocity across the electrode array (**Figure 1C**). As previously described, the spike amplitude is an indicator of depolarization (akin to an electrocardiogram QRS complex) and FPD is an indicator of repolarization (akin to QT interval)(26). Contractility parameters of interest included excitation contraction delay and contraction beat amplitude (**Figure 1D**). Excitation contraction delay quantifies the time between the depolarization spike and the contractility waveform peak, while the beat amplitude is measured by a change in impedance as hiPSC-CMs contract and relax (40, 41). In a subset of experiments, cardiomyocyte-electrode coupling was enhanced to record local extracellular action potentials (LEAP) (28) to calculate the action potential (APD) duration at 30% (APD30), 50% (APD50), and 90% (APD90) repolarization (**Figure 1E**). Data points from individual wells were excluded if hiPSC-CMs were not beating at the desired pacing frequency, or if the signal-to-noise ratio was too small for accurate measurements.

**Figure 1.**
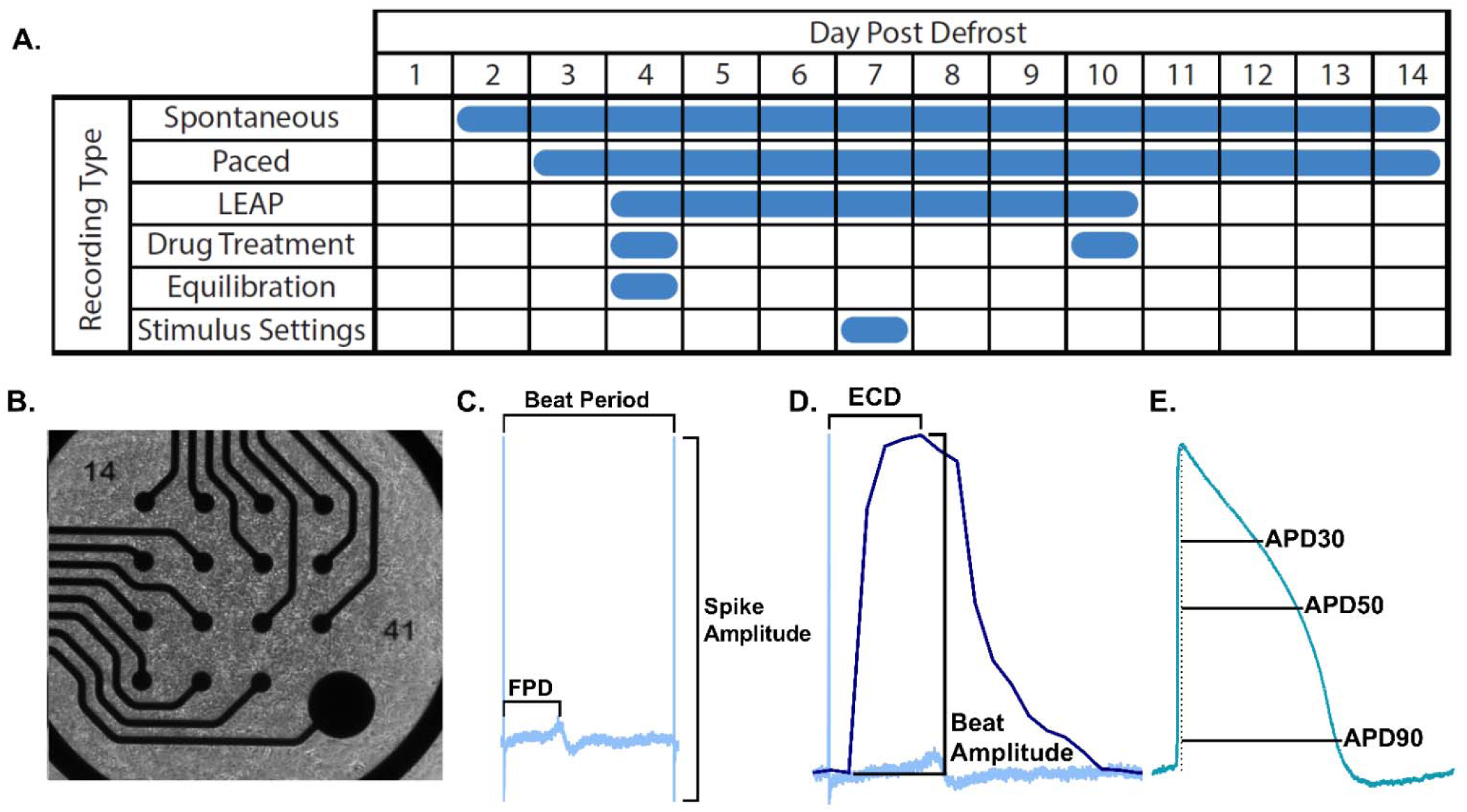
Experimental Design. **A**) Timeline of microelectrode array (MEA) recordings. **B)** Example image of human induced pluripotent stem cell derived cardiomyocytes (hiPSC-CMs) plated onto an MEA. **C)** Representative extracellular field potential waveform, with markers indicating the beat period, field potential duration (FPD), and spike amplitude. **D)** Representative contraction recording, with markers indicating the excitation-contraction delay (ECD) and beat amplitude. **E)** Representative local extracellular action potential recording with markers indicating action potential duration (APD) at 30% (APD30), 50% (APD50), and 90% (APD90) repolarization.

*Daily hiPSC-CM maturation:* We tested the impact of cell culture duration on hiPSC-CM physiology by acquiring biosignals each day during the first two weeks after defrost (**Figure 1A**). Field potential recordings and contractility measurements were acquired from spontaneously beating hiPSC-CMs 2 to 14 days post-defrost; these signals were also recorded in response to external pacing (3 to 14 days post-defrost) after cells were fully attached and formed a confluent monolayer. LEAP recordings were also acquired in response to external pacing (4 to 10 days post-defrost). For paced recordings, hiPSC-CMs were stimulated at 1.5 Hz (stimulus settings: 400 µs, 800 mV, and 20 µA).

*Electrical stimulation settings*: We tested the impact of adjusting stimulation settings on cardiac electrophysiology endpoints. To negate the impact of cell culture time, these experiments were only performed on day 7 post-defrost. Variations in current (20-50 µA), stimulus duration (400-1000 µs), and voltage (800-1200 mV) were tested. Only one parameter was adjusted at a time; otherwise, the settings were held constant (stimulus duration: 400 µs; voltage: 800 mV; current: 20 µA).

*Drug Response:* To measure the impact of cell culture time on hiPSC-CM drug responsiveness, cells were exposed to three different drugs at two different time points (day 4, day 10 post-defrost; **Figure 1A**). A hERG channel blocker (50 nM E-4031, #15203 Cayman Chemical) was tested for its ability to prolong FPD and APD(28, 42, 43). E-4031 reportedly suppresses spontaneous activity at concentrations above 30 nM (44). A calcium channel blocker (100 nM nifedipine, #N7634-1G Sigma-Aldrich) was tested for its ability to shorten APD (28) and increase the spontaneous beating rate (43). A beta-adrenergic agonist (100 nM isoproterenol, #I5627-25G; Sigma-Aldrich) was tested for its ability to shorten APD, increase the spontaneous beating rate, and increase conduction velocity(44, 45). Since these pharmacological agents can alter the spontaneous beating rate, electrical signals were recorded in response to external pacing (1.5 Hz for E-4031, 2-3 Hz for nifedipine and isoproterenol). Stimulus settings were the same for all drug exposure studies (stimulus duration: 400 µs; voltage: 800 mV; current: 20 µA). Drug treatments were applied for 15-minutes before recording electrical signals.

### Data acquisition and statistical analysis

For each measurement, biosignals were collected for 30 seconds to 2 minutes using an MEA system (12.5 kHz sampling frequency). Each MEA file was then manually reviewed (DG, JP) to select one electrode per well with the largest signal amplitude (e.g., action potential or t-wave). Whenever possible, signals were selected for FPD analysis that had an upright t-wave. After manually selecting one electrode per well, waveforms from that recording electrode were averaged (20 seconds). Individual wells were excluded if the signal amplitude was insufficient for accurate measurements, or when hiPSC-CMs were not beating at the desired pacing frequency. Outliers were identified and removed using the ROUT method (5% false discovery rate). Results are reported throughout as mean ± standard deviation. Statistical analysis was performed using either a Student’s t-test (two groups), one– or two-way analysis of variance (ANOVA) with Holm-Sidak multiple comparisons test (three or more groups), as indicated in each figure legend. An adjusted p-value <0.05 was considered statistically significant. Data are graphically represented as the mean with range (min to max), unless otherwise indicated. Replicates are defined in each figure.

## Results

### MEA equilibration time impacts hiPSC-CM electrophysiology measurements

We used a commercially available hiPSC-CM cell line (#01434, Fujifilm Cellular Dynamics) that has been widely used in preclinical cardiac safety pharmacology studies (27, 30–32, 46–49). The manufacturer’s instructions (50) recommend equilibrating hiPSC-CMs in an MEA system for approximately 10 minutes before recording electrical signals, although some studies have reported using a longer equilibration time (≥15-20 minutes)(43, 51–53). To determine whether the equilibration period influenced experimental results, we continuously recorded electrical signals after transferring hiPSC-CM culture plates to an MEA system. Over the course of a 20-minute recording, the spontaneous beating rate increased markedly by 22.6% (*2 minutes*: 40.2±4.8, *20 minutes:* 49.3±5.9 BPM, p<0.0001; **Figure 2A**) and the FPD shortened by 7.7% (*2 minutes:* 385.8±19.8, *20 minutes:* 356.1±18.8 ms, p<0.0001; **Figure 2C-D**). Adaptations in the spontaneous beating rate and FPD were most exaggerated at the beginning of the recording, reaching a plateau near the end of the equilibration period. For example, compared to the last measurement at 20-minutes, the spontaneous beating rate was significantly slower only between 2-12 minutes (p<0.05) and FPD was longer only between 2-7 minutes (p<0.05). FPD plateaued as the cells reached a steady stated, with a mean difference of less than 6 ms for each measurement between 8-20 minutes. No significant differences were observed in the depolarization spike amplitude or conduction velocity during the equilibration period (**Figure 2E-H**). To reduce hiPSC-CM equilibration as a confounding factor, it is recommended that hiPSC-CM acclimate to the MEA system for 10-20 minutes before performing subsequent experiments. For reference, continuous measurements (in 60 second intervals) are available in **Supplemental Table 1**.

**Figure 2.**
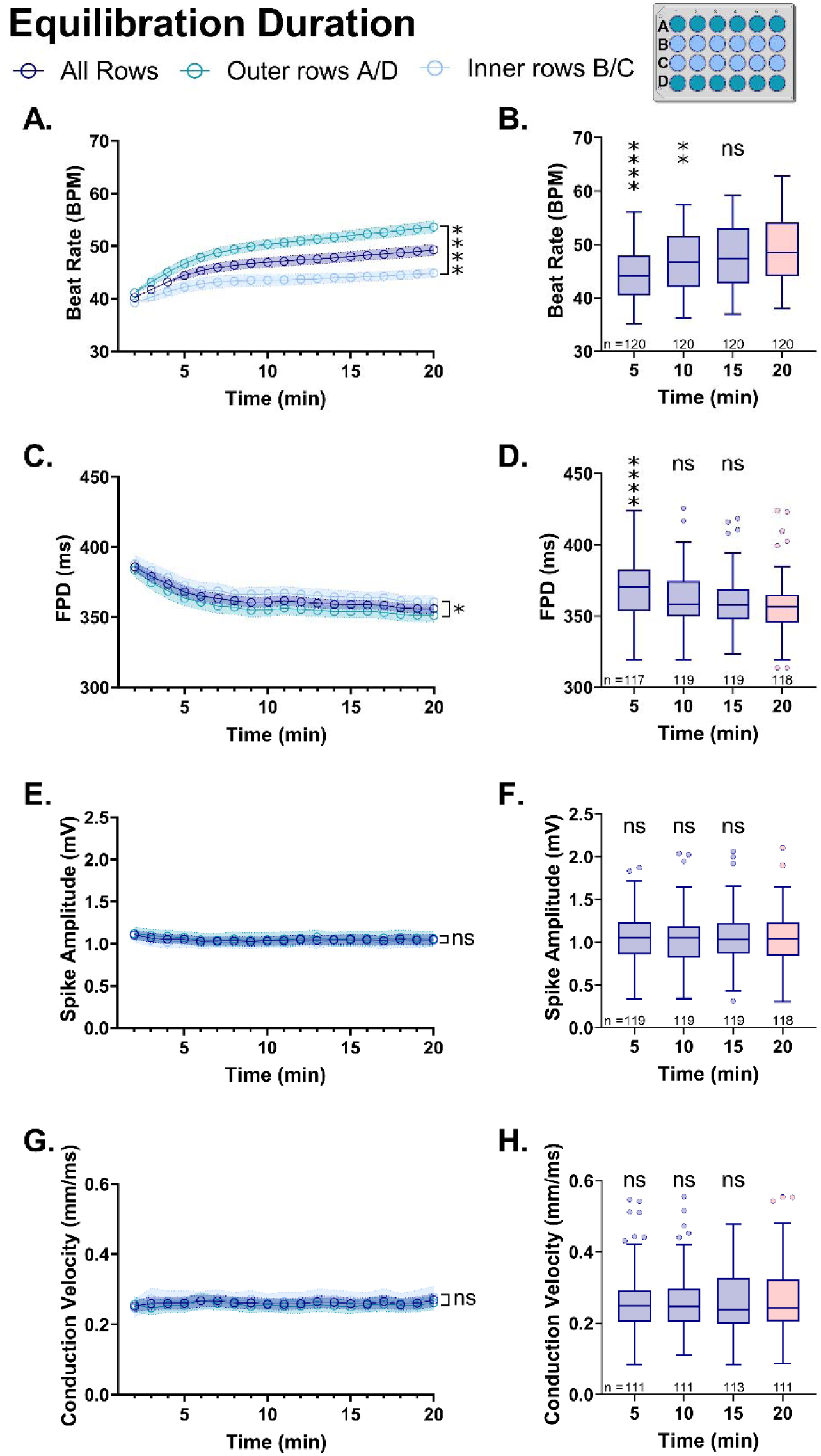
Equilibration Duration. hiPSC-CM electrophysiology data was collected continuously for 20 minutes (4 days post-defrost) during spontaneous beating. **A,B)** Spontaneous beating rate. **C,D)** Field potential duration (FPD). **E,F)** Depolarization spike amplitude. **G,H)** Conduction velocity. **Left:** Row location measurements are reported as the average of all wells across the MEA plate (purple), outer rows (green), or inner rows (blue). Open circle = mean, shaded region = 95% confidence interval. Statistical comparison was performed at 20-minute timepoint (ANOVA with Holm-Sidak correction for multiple comparisons). All Rows: n=107-120; Outer rows A/D: n=54-60; inner rows B/C: n=51-60. **Right:** Equilibration time measurements are for all wells shown as a Tukey box-and-whisker plot, with outliers indicated. Statistical comparisons at five-minute intervals (purple box plots) versus the end of equilibration at 20-minute timepoint (red box plot). ANOVA with Holm-Sidak correction for multiple comparisons. *p<0.05, **p<0.01, ****p<0.0001. Number of replicates is indicated for each time point.

Unexpectedly, we identified location specific differences in hiPSC-CM electrophysiology measurements according to the well location on the MEA plate (**Figure 2A-D**). Throughout the 20-minute equilibration period, we noted a significantly faster beating rate in the outer rows (“A” and “D”) and inner rows (“B” and “C”), which became exaggerated over time. At the 2 minute recording time, a modest 4.8% difference was observed between outer (41.1±2.3 BPM) and inner rows (39.2±2.8 BPM, p<0.001), which increased to a 16.3% difference at the 20 minute recording time (*outer rows:* 53.7±4.5, *inner rows*: 44.9±3.4 BPM, p<0.0001; **Figure 2A**). FPD measurements also differed by well location; at the 20 minute recording time hiPSC-CMs plated in the outer rows had a shorter FPD (351.2±18.0 ms) compared to cells in the inner rows (360.9±18.5 ms, p<0.05; **Figure 2C**). No significant difference in spike amplitude or conduction velocity was observed between the inner and outer rows of hiPSC-CM (**Figure 2E,G**). To reduce well location as a confounding factor, it is recommended that hiPSC-CM treatment groups are dispersed across the MEA multiwell plate – rather than grouping a specific treatment group across a single row or column.

### Electrical stimulation settings influence hiPSC-CM electrophysiology measurements

Empirically, we noted that certain pharmacological agents altered the excitability of hiPSC-CMs, preventing the cell monolayer from capturing at a desired pacing frequency. This finding has been reported by others, wherein hiPSC-CMs continue to beat spontaneously at ½ or ⅓ the applied pacing rate (54). To remedy this problem, it can be helpful to adjust the electrical stimulation settings (i.e., pulse duration, voltage amplitude, and/or current) in order to successfully pace hiPSC-CMs at a desired frequency. We measured hiPSC-CM electrophysiology endpoints in response to standard electrical stimulations settings, and then modified each variable independently to quantify changes in the results (repeated-measures design, **Figure 3**). FPD measurements were slightly shorter when the current was increased from 20 to 50 μA (308.9±12 to 304.9±1.7 ms, p<0.001), but unchanged by adjustments in the pulse duration or voltage amplitude (**Figure 3A-C**). Spike amplitude and conduction velocity measurements were influenced by changes in current, duration, and voltage amplitude (**Figure 3D-I**). For example, a faster conduction velocity was noted when the pulse duration was lengthened from 400 to 1000 μs (0.207±0.02 to 0.215±0.03 mm/ms, p<0.001). To avoid misinterpreting results that are influenced by electrical stimulation settings, we recommend using a predefined setting to pace hiPSC-CMs at multiple frequencies (1-3 Hz). Such an approach will allow for a direct comparison of rate-dependent electrophysiology metrics before and after drug/chemical treatment, particularly in cases where cell excitability or the intrinsic beating rate changes.

**Figure 3.**
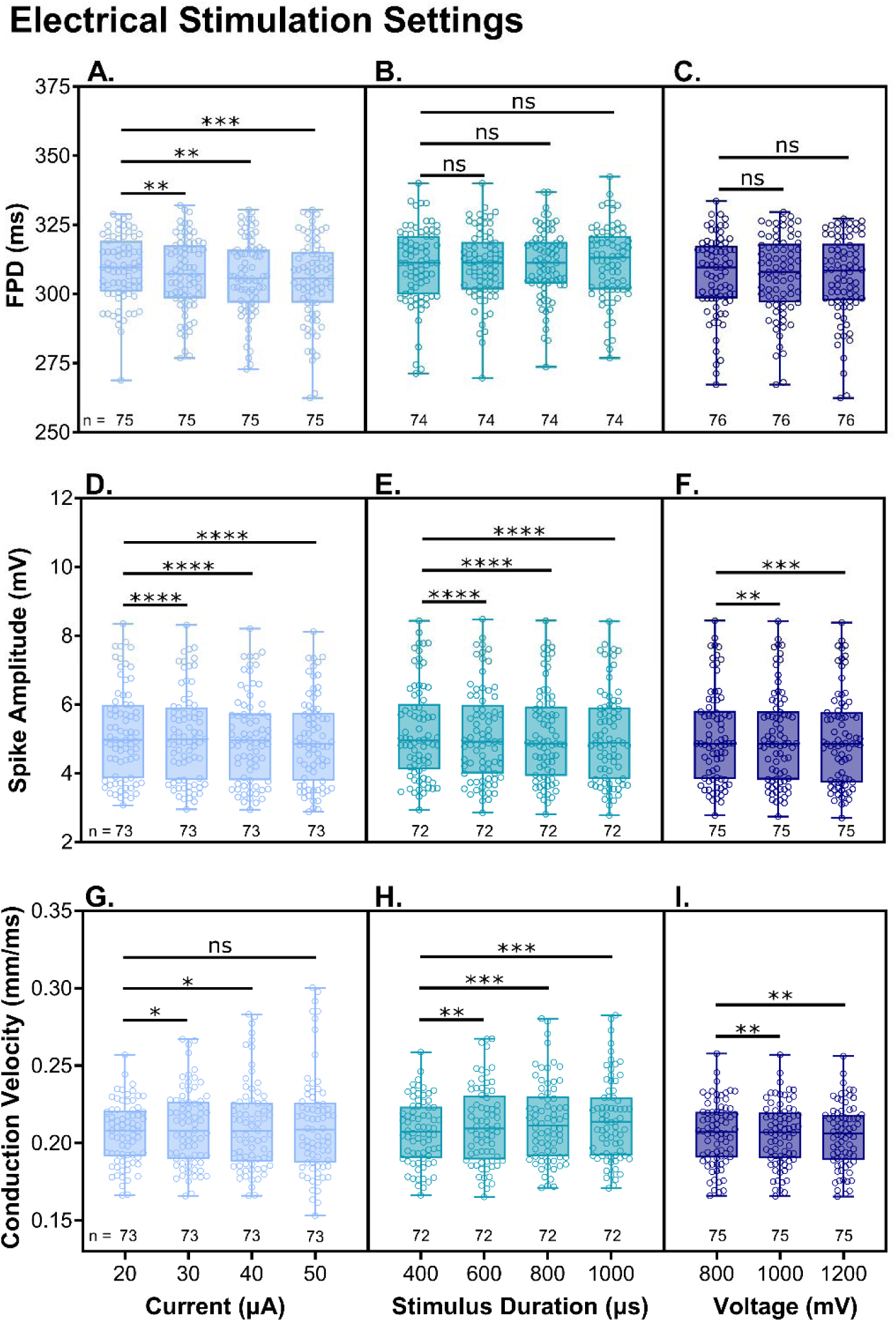
Electrical stimulation settings. hiPSC-CM electrophysiology data was collected (7 days post-defrost) in response to external pacing at 1.5 Hz frequency (standard settings: 400 µs stimulus duration, 800 mV voltage amplitude, 20 µA current). Each parameter was adjusted independently, with statistical comparison versus the standard default setting. **Top:** Changes in field potential duration (FPD) measurement following adjustment in **A)** applied current, **B)** stimulus duration, **C)** voltage amplitude. **Middle:** Changes in spike amplitude following adjustment in **D)** applied current, **E)** stimulus duration, **F)** voltage amplitude. **Bottom:** Changes in conduction velocity following adjustment in **G)** applied current, **H)** stimulus duration, **I)** voltage amplitude. Measurements are shown as a box-and-whisker plot (all values shown, min to max). Statistical comparison versus default standard setting. ANOVA with Holm-Sidak correction for multiple comparisons. *p<0.05, **p<0.01, ***p<0.001, ****p<0.0001. Number of replicates indicated.

### Changes in hiPSC-CM electrophysiology during cell culture

Previous studies have demonstrated that stem cell-derived cardiomyocytes continue to mature with longer cell culture time, which can directly impact cardiomyocyte electrophysiology(18, 55–57). Preclinical drug testing studies frequently use commercially available hiPSC-CM cell lines within the first 2 weeks post-defrost. Therefore, we characterized daily phenotypic changes in iCell cardiomyocytes^2^ within this timeframe while cells were spontaneously beating (**Supplemental Table 2**). The spontaneous beating rate decreased 23.6% within the first 14 days of cell culture (*2 days:* 53.8±5.1, *7 days:* 46.2±7.8, *14 days:* 41.1±4.5 BPM; p<0.0001; **Figure 4A**). Per the manufacturer’s instructions, the ‘optimal’ day for hiPSC-CM experimentation is 7 days post-defrost (50). Compared to this 7-day timepoint, the beating rate was significantly different 2-5 days and 9-14 days post-defrost (p<0.001) – only 6 and 8 days post-defrost were comparable (**Figure 4A**). We also observed a 1100% increase in the depolarization spike amplitude and 90% lengthening of FPD with longer cell culture time, which was exaggerated during the first week and plateaued slightly during the second week post-defrost (**Figure 4C-F**). For example, at 2 days post-defrost the mean FPD was 288.9±15.7 ms, which increased to 450.6±39.5 ms at 7 days and 545.9±24.2 ms at 14 days. The depolarization spike amplitude started at 0.30±0.24 mV at 2 days post-defrost and increased drastically to 2.45±0.76 mV at 7 days and 3.61±0.77 mV at 14 days. Conduction velocity exhibited a sinusoidal trend across the first two weeks of culture with an initial decrease in conduction velocity from 2-5 days post-defrost (*2 days*: 0.33 ± 0.09, *5 days*: 0.21 ± 0.05 mm/ms), followed by a gradual increase (*7 days*: 0.26 ± 0.06 mm/ms), and then another decline from 10-14 days (*10 days*: 0.35 ± 0.12 mm/ms, *14 days*: 0.30 ± 0.06 mm/ms; **Figure 4G-H**).

**Figure 4.**
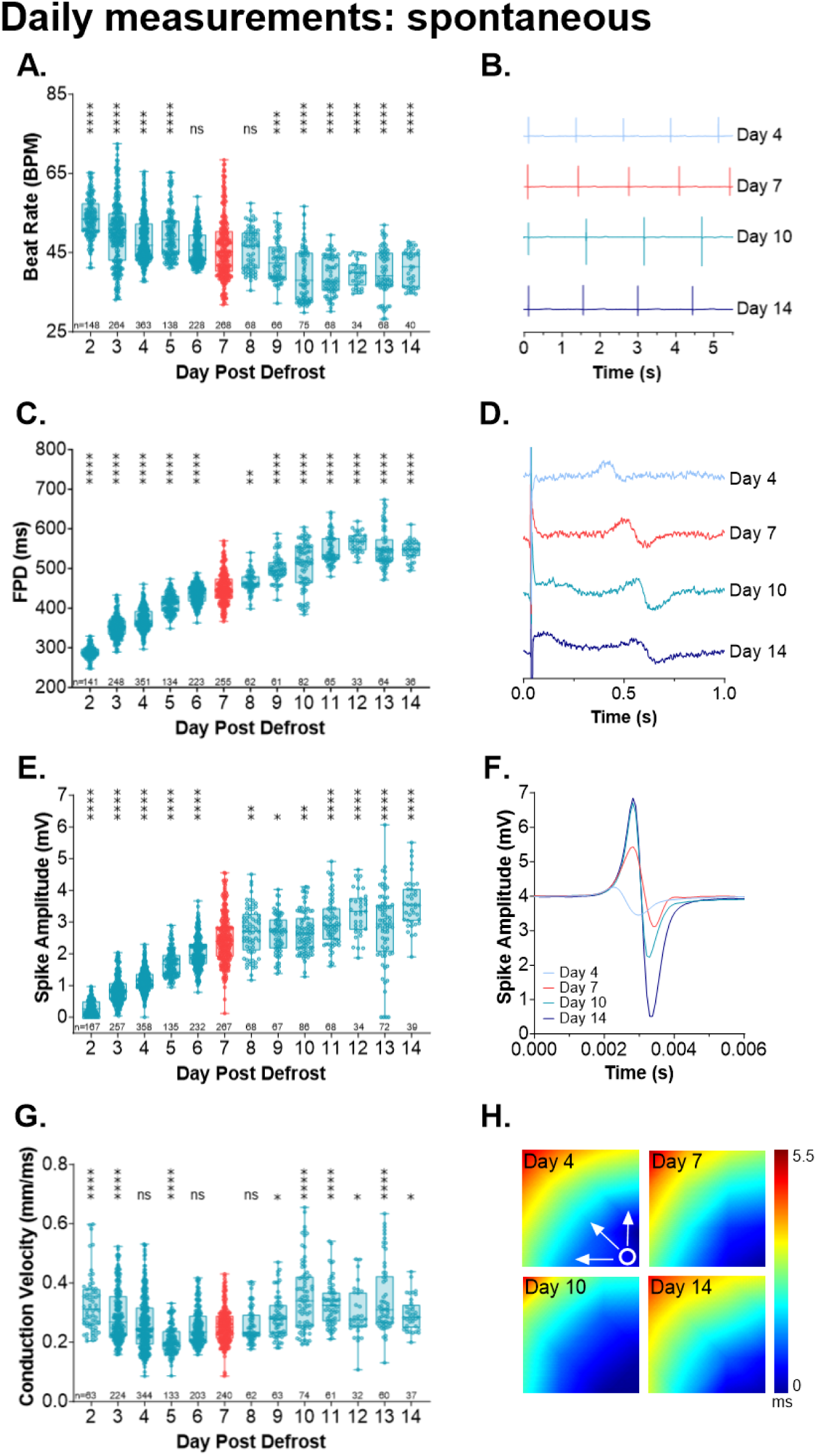
Daily changes in hiPSC-CM electrophysiology measurements (spontaneously beating). During two weeks of cell culture, **A,B)** spontaneous beating rate slows, **C,D)** field potential duration (FPD) lengthens, and **E,F)** depolarization spike amplitude increases. **G)** Conduction velocity changed in a sinusoidal manner during two weeks of cell culture. **H)** Activation maps show electrical signal propagating across a cardiomyocyte layer at different timepoints. **Left:** Daily measurements are shown as a box-and-whisker plot (all values shown, min to max), with statistical comparison versus day 7 values (shown in red). ANOVA with Holm-Sidak correction for multiple comparisons. *p<0.05, **p<0.01, ***p<0.001, ****p<0.0001. Number of replicates indicated. **Right:** Representative traces at different timepoints.

Since the hiPSC-CM spontaneous beating rate slowed significantly during 2-weeks of cell culture and cardiac electrophysiology metrics are rate-dependent, we measured each variable again in response to external pacing (1.5 Hz). In response to pacing, FPD measurements were longer in hiPSC-CMs that were cultured for longer periods of time – although the magnitude of this change (21.9%; *3 days:* 275.5±13.8, *14 days*: 336.0±14 BPM, p<0.0001) was less pronounced that observed during spontaneous beating (54.8% *3 days vs 14 days*). Changes in paced FPD values were exaggerated during the first week of culture, and then plateaued after 9 days in culture (**Figure 5A**). In response to pacing, the depolarization spike amplitude also increased with longer cell culture time (44.4%; *3 days*: 3.8±1.1, *14 days*: 6.9±0.9 mV, p<0.0001; **Figure 5B**) – but again, the magnitude was less pronounced than observed during spontaneous beating (76.2%; *3 days vs 14 days*). The depolarization spike amplitude was significantly larger when cells were paced versus spontaneously beating, for every time point investigated (e.g., *7 days spontaneous:* 2.5±0.8 mV, *7 days paced:* 5.4±1.3 mV). Paced conduction velocity measurements exhibited a sinusoidal trend across the first two weeks of culture, initially decreasing from 3-5 days post-defrost (3 *days:* 0.24±0.02, *5 days:* 0.20±0.02 mm/ms), then increasing during the remainder of the study period (*7 days*: 0.23±0.02, *14 days*: 0.28±0.02 mm/ms; **Figure 5C**). For reference, daily measurements are available in **Supplemental Table 3**.

**Figure 5.**
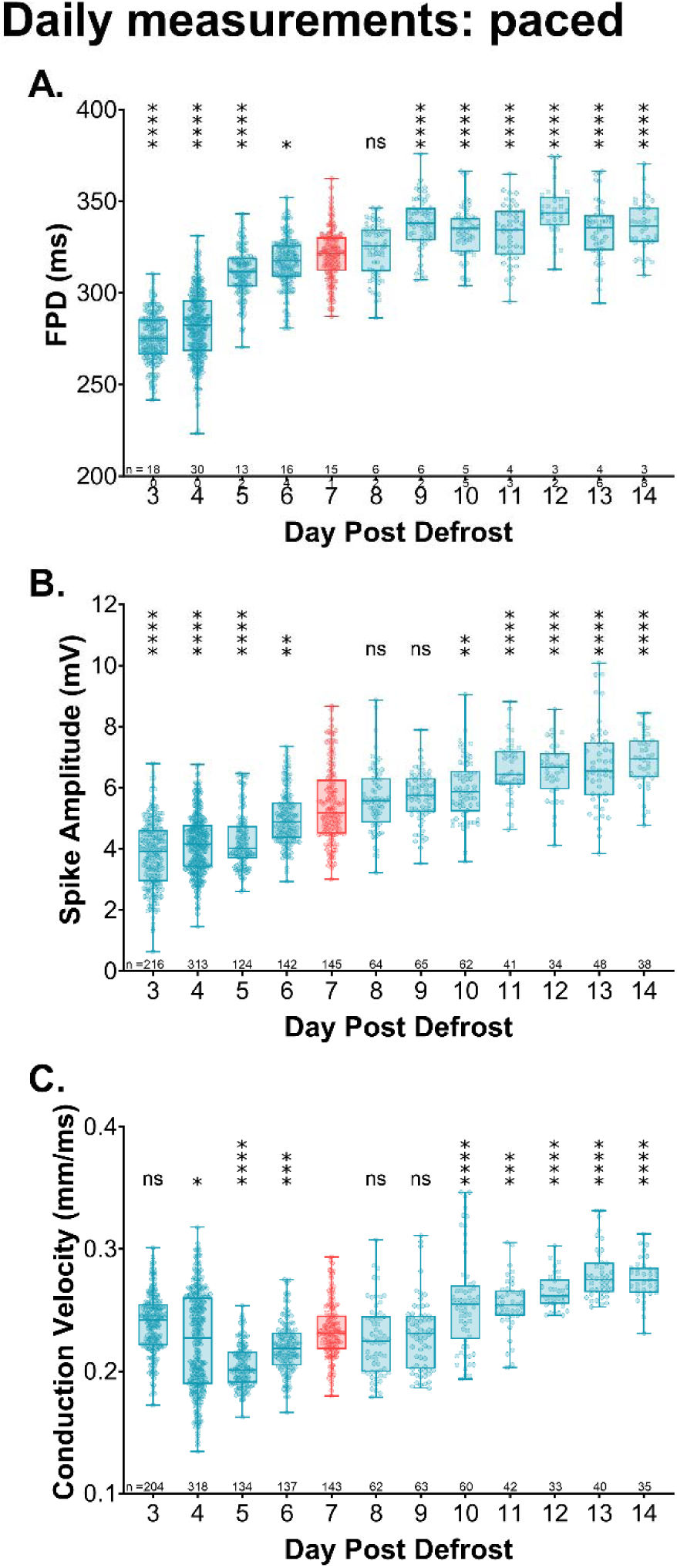
Daily changes in hiPSC-CM electrophysiology measurements (external pacing, 1.5 Hz). During two weeks of cell culture, **A)** field potential duration (FPD) lengthens, **B)** depolarization spike amplitude increases, and **C)** conduction velocity changes in a sinusoidal manner. Daily measurements are shown as a box-and-whisker plot (all values shown, min to max), with statistical comparison versus day 7 values (shown in red). ANOVA with Holm-Sidak correction for multiple comparisons. *p<0.05, **p<0.01, ***p<0.001, ****p<0.0001. Number of replicates indicated.

MEA technology allows for the measurement of both extracellular field potentials (akin to the electrocardiographic QT interval) and local extracellular action potentials (akin to the cardiac action potential). It has previously been documented that the FPD measurement strongly correlates with an APD between 50-90% repolarization (28, 58). Similar to our FPD measurements, we found that the APD at 30% repolarization lengthened by 18.6% across multiple days in cell culture (*4 days:* 171.7±19.4, *10 days*: 203.7±26.0 ms, p<0.01; **Figure 6A**).

**Figure 6.**
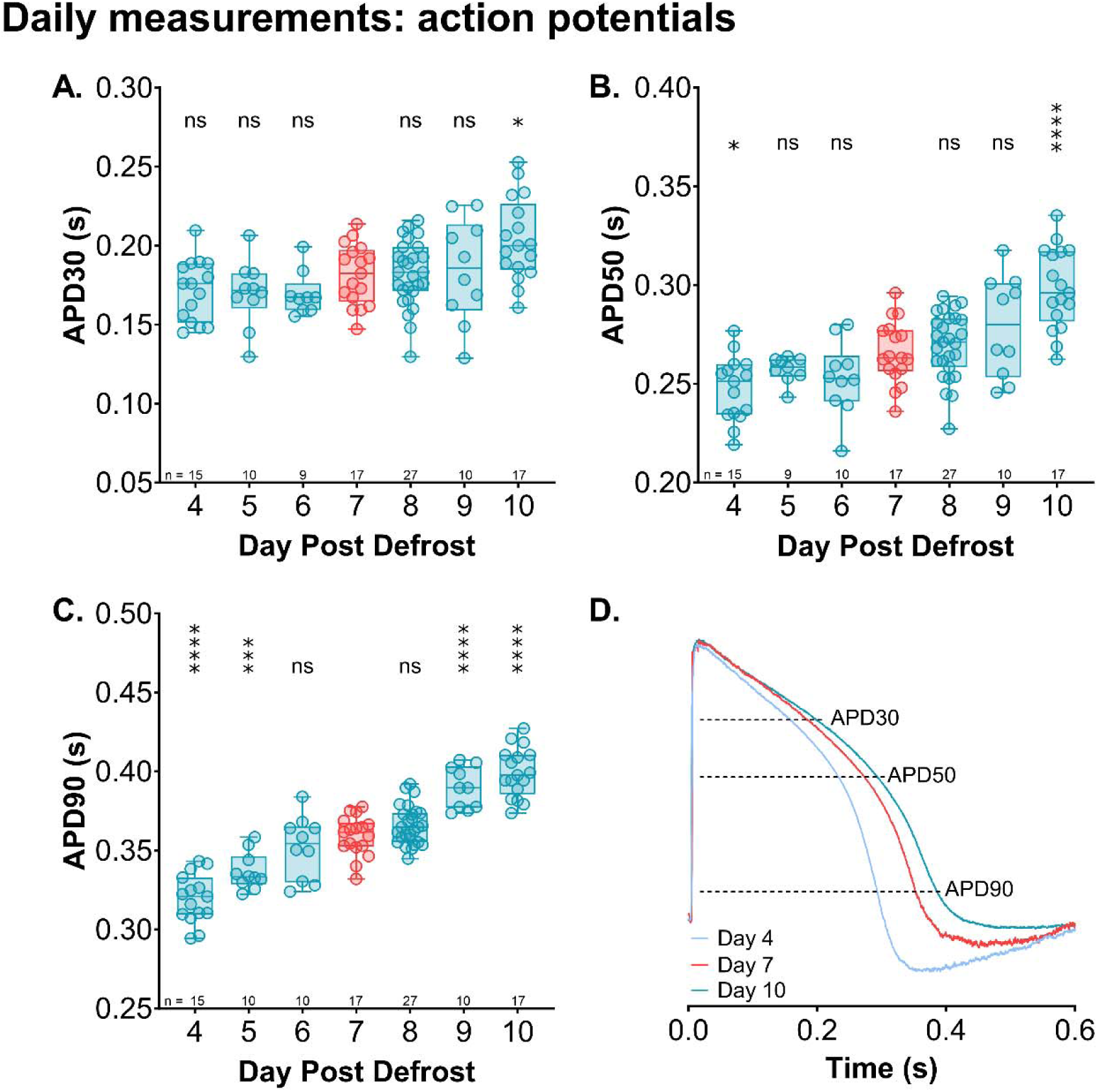
Daily changes in hiPSC-CM action potential duration (external pacing, 1.5 Hz). **A**) action potential duration at 30% repolarization (APD30), **B)** 50% repolarization (APD50), **C)** 90% repolarization (APD90). **D)** Representative example of action potential recordings. Daily measurements are shown as a box– and-whisker plot (all values shown, min to max), with statistical comparison versus day 7 values (shown in red). ANOVA with Holm-Sidak correction for multiple comparisons. *p<0.05, ***p<0.001, ****p<0.0001. Number of replicates indicated.

APD lengthening was more dramatic at later phases of repolarization, increasing by 20.5% at APD50 (*4 days:* 247.6±16.4, *10 days*: 298.4±20.5 ms, p<0.01; **Figure 6B**) and 24.9% at APD90 (*4 days:* 319.6±15.3, *10 days*: 399.2±15.5 ms, p<0.01; **Figure 6C**). When compared to the ‘optimal’ day for hiPSC-CM experimentation (7 days post-defrost (50)), only 6-8 days post-defrost produced comparable APD90 values. For reference, daily measurements are available in **Supplemental Table 4**. We recommend using hiPSC-CM within a narrow timeframe after defrost (e.g., 6-8 days), and reporting the raw values for each cardiac metric to facilitate data interpretation and comparison between research laboratories.

### Stability of hiPSC-CM contractility parameters during cell culture

Previous studies have demonstrated that hiPSC-CMs continue to mature in cell culture. Younger hiPSC-CMs have longer contractile events with shorter calcium transient amplitudes, while older cells display faster contraction, taller calcium transient amplitudes, and prolonged relaxation (4 weeks)(56, 57). In the current study, we only observed slightly differences in excitation-contraction delay and the contraction beat amplitude – and these adaptations were limited to the first few days of cell culture as the monolayer was forming (**Figure 7**). The time delay between excitation and contraction lengthened from 2-4 days (*2 days*: 248.8±30.6, *4 days*: 277.5±29.5, p<0.0001; **Figure 7A**) and then plateaued, but only during spontaneous beating. The contraction beat amplitude beat amplitude decreased in the first few days of cell culture, during both spontaneous (*2 days*: 2.2±0.4, *5 days*: 1.7±0.3%) and paced (*3 days*: 1.6±0.3, *5 days:* 1.4±0.3%) recordings (**Figure 7B,D**). Since these experiments were only performed up to 14 days in culture, it is possible that additional adaptations occur at later timepoints.

**Figure 7:**
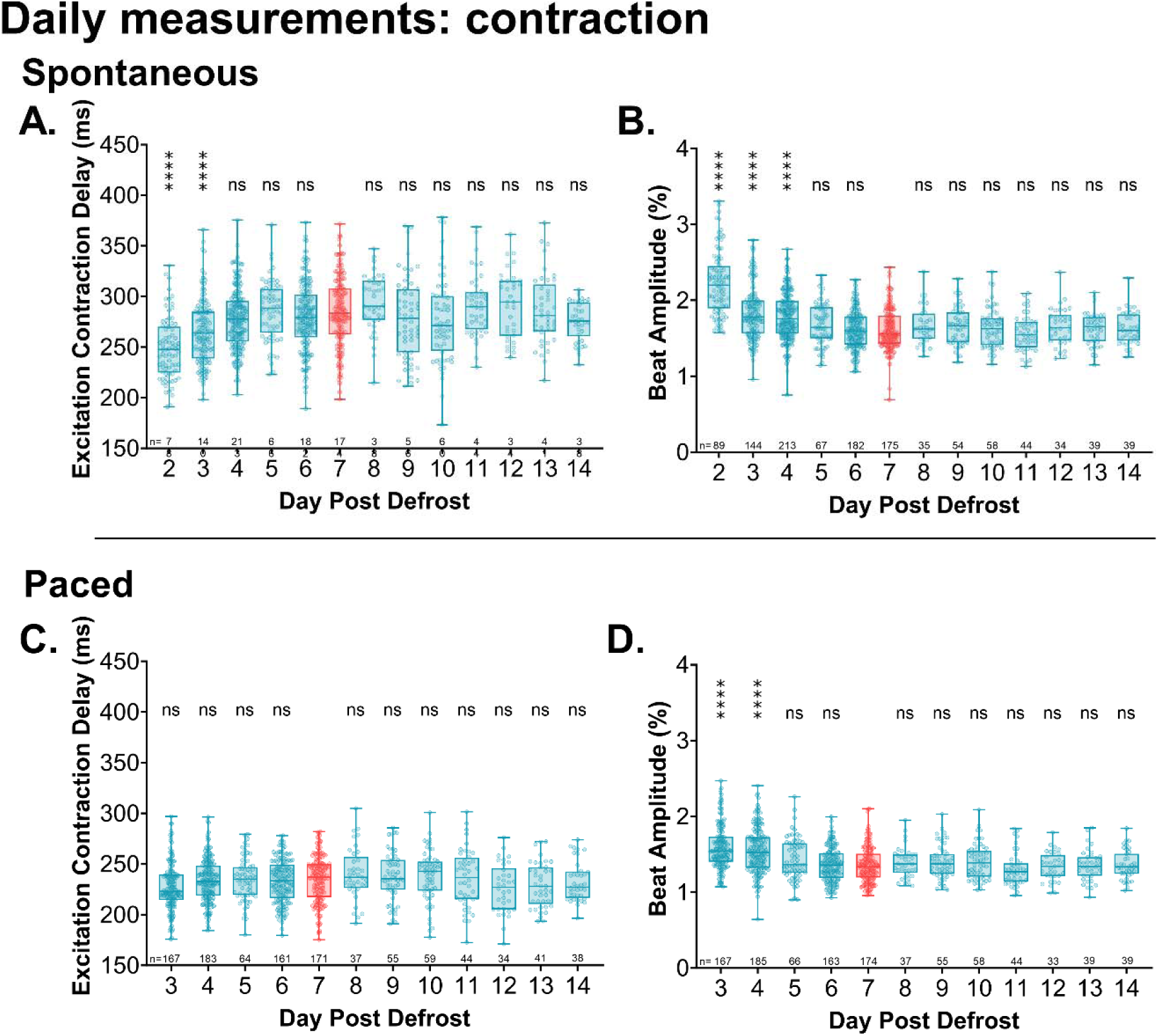
Daily changes in hiPSC-CM contraction measurements. **A,C**) Daily changes in excitation-contraction delay. **B,D)** Daily changes in beat amplitude. **Top:** Measurements collected daily during two weeks of cell culture, while cells were spontaneously beating. **Bottom:** Measurements in response to external pacing (1.5 Hz). Daily measurements are shown as a box-and-whisker plot (all values shown, min to max), with statistical comparison versus day 7 values (shown in red). ANOVA with Holm-Sidak correction for multiple comparisons. *p<0.05, **p<0.01, ***p<0.001, ****p<0.0001. Number of replicates indicated.

### Changes in hiPSC-CM drug responsiveness during cell culture

Previous studies have reported profound differences in cardiac gene expression and key ionic currents in stem cell-derived cardiomyocytes after long-term cell culture (weeks to months)(18, 56, 57) and/or the implementation of maturation-specific protocols (16–21). We investigated whether such underlying developmental differences could alter drug responsiveness, even within the relatively short-time frame recommended for hiPSC-CM experimentation (50). First, we observed that the spontaneous beating rate was altered by the application of a β-adrenergic agonist (isoproterenol), calcium channel blocker (nifedipine), or hERG blocker (E-4031) – but the magnitude of this change was influenced by cell culture duration (**Figure 8A**). Isoproterenol caused a slight increase in the beating rate (11.5%) on day 4 post-defrost, which was more pronounced (48.8%) on day 10. Conversely, nifedipine increased the beating rate (112.9%) on day 4 post-defrost, but its effect was more tempered (56.8%) on day 10. E-4031 caused a slight increase in the beating rate (4.5%) on day 4 post-defrost, while significant beat rate slowing was observed (–19.1%) on day 10 post-defrost. Second, the application of E-4031 lengthened the FPD (*4 days:* 31.2%, *10* days: 17.2%) and APD (*4 days*: 41.5%, *10 days*: 10.0%; **Figure 8C,H**) but the effects were more exaggerated at the earlier timepoint. Third, the application of isoproterenol increased conduction velocity at the later timepoint (*4 days:* –6.5%, *10 days*: 3.5%) and the depolarization spike amplitude was also more affected at the later timepoint (*4 days:* –12.3%, *10 days*: –27.3%; **Figure 8D,E**). The length of cell culture time should be considered when performing drug screening or safety testing, as hiPSC-CMs may respond differently depending on their maturation state.

**Figure 8:**
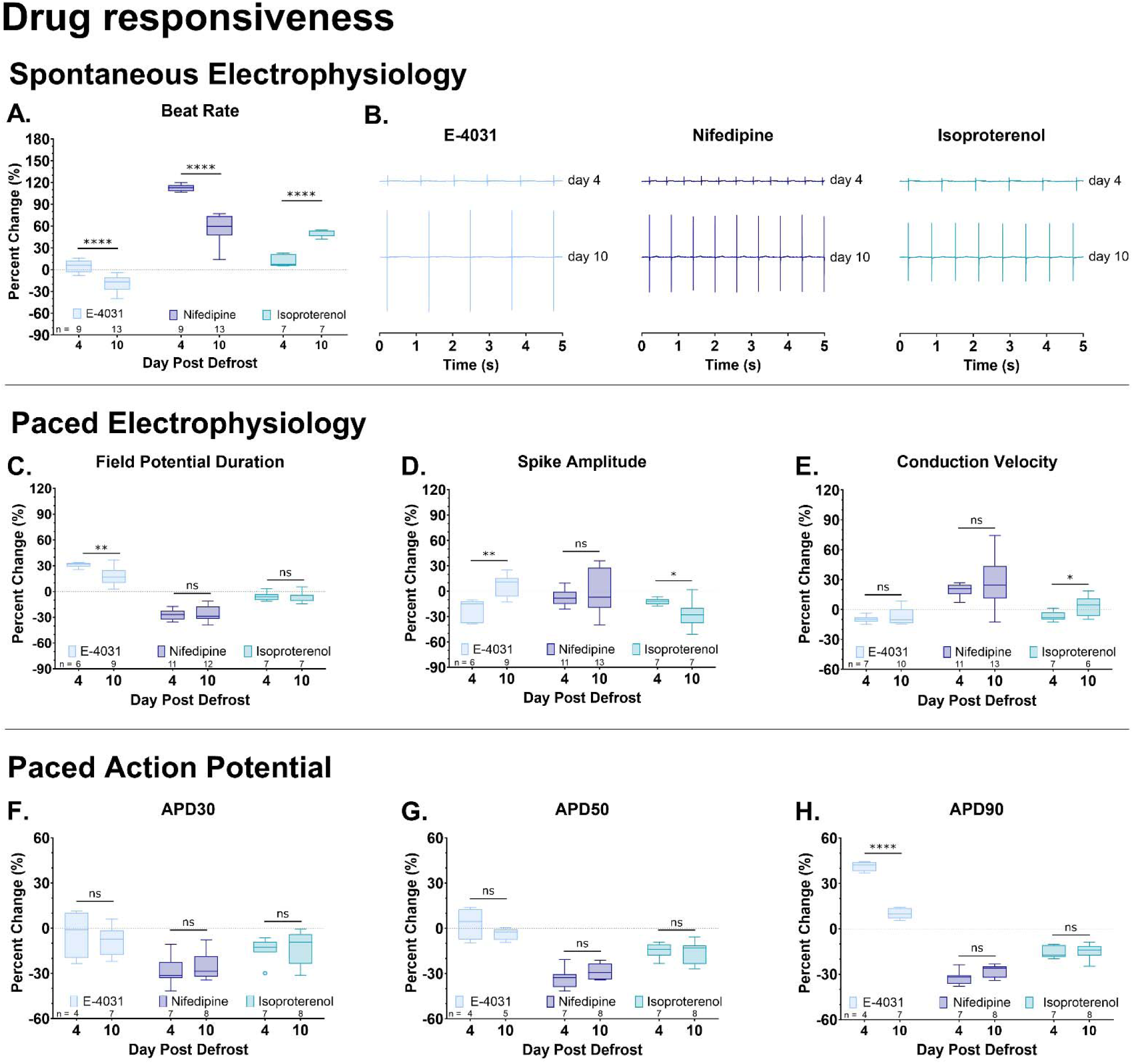
hiPSC-CM drug responsiveness changes based on the length of cell culture time. hiPSC-CM electrophysiology data was collected (4 vs 10 days post-defrost) at baseline and after exposure to a hERG channel blocker (E-4031), calcium channel blocker (nifedipine), or beta-adrenergic agonist (isoproterenol). **A)** Percent change in the spontaneous beating rate, normalized to baseline measurement (repeated-measures) after 15-minute drug treatment. **B)** Representative traces of spontaneous beating after drug treatment. **C-E)** Percent change in the field potential duration, spike amplitude, and conduction velocity (normalized to baseline measurement) after 15-minute drug treatment. hiPSC-CMs were externally paced to normalize the beating rate (1.5 Hz for E-4031 which slowed the intrinsic rate; 3 Hz for nifedipine and isoproterenol which increased the intrinsic rate). **F-H)** Percent change in the action potential duration at 30%, 50%, and 90% repolarization (normalized to baseline measurement) after 15-minute drug treatment. hiPSC-CMs were externally paced to normalize the beating rate (1.5 Hz for E-4031; 2 Hz for nifedipine and isoproterenol). Measurements are shown as a Tukey box-and-whisker plot. Unpaired Student’s t-test, two tailed. *p<0.05, **p<0.01, ****p<0.0001. Number of replicates indicated.

## Discussion

In the presented study, we utilized a commercially available hiPSC-CM cell line (#01434, Fujifilm Cellular Dynamics) that has been widely used to test the efficacy, safety, and toxicity of pharmacological agents and environmental chemicals (47, 59–62). To the best of our knowledge, this is the first study to monitor daily changes in hiPSC-CM physiology throughout 14-days in cell culture, which is within the manufacturer’s recommended timeframe for experimentation. Our study identified multiple sources of variability that impacted hiPSC-CM MEA measurements, including: 1) well-to-well variability across the MEA plate, 2) length of equilibration period as hiPSC-CMs reach a steady state within the MEA system, 3) electrical stimulation settings used to externally pace hiPSC-CMs, and 4) duration of post-defrost cell culture. Collectively, our study highlights important user-defined experimental factors that should be considered when using hiPSC-CM models for disease modeling, safety pharmacology, and cardiotoxicity screening.

In our study, we were surprised to find that the well location within an MEA plate impacted baseline electrophysiology measurements. Specifically, hiPSC-CMs plated in the top/bottom rows had faster beating rates and shorter FPD values compared to hiPSC-CMs plated in the inner rows. These results highlight an important consideration, as the location of treatment groups on an MEA plate could give the (mis)perception of drug effects (e.g., vehicle control treatment added only to the top row of cells, drug treatment in a middle row). Potential explanations for this variability may include slight differences in temperature, pH, or CO_2_ distribution across the plate. Unfortunately, we were not able to pinpoint the specific cause of these differences, since measuring the cell culture media temperature or gas concentration required opening the MEA system (which in turn, further disturbs the atmosphere). Well-to-well variability persisted even after allowing hiPSC-CMs to equilibrate to the new environment within the MEA system for 20-minutes. Further, we noted a progressive increase in beating rate and shortening of FPD during this equilibration period. Accordingly, even slight modifications in equilibration time between studies can influence hiPSC-CM MEA data and may contribute to inter-facility variability. For example, Nozaki et al. reported that hiPSC-CM MEA studies accurately predict the risk of drug-induced arrhythmias – but the minimum concentration required to produce arrhythmia-like waveforms can vary by 10-fold when results are compared between multiple laboratories (despite using the same hiPSC-CM cell line and batch) (34). Kitaguchi et al. also documented intra-facility and inter-facility variability in basal hiPSC-CM measurements and drug response, and reported that the drug concentration inducing FPD prolongation ranged from 1.8-5.8-fold between different laboratories using the same cell line (63). As another example, Blinova et al. reported inter-facility variability between 10 different laboratories (0.69-0.88 cross-site correlation) in detecting drug-induced repolarization prolongation in 28 tested drugs (30). Potential sources of such variability may include well-to-well differences, slight deviations in equilibration time, minor modifications in cell culture techniques (e.g., plating density, age of cells) or MEA recording parameters. Indeed, in this study we found that minor adjustments in the pulse width, voltage amplitude, or applied current can alter cardiac electrophysiology measurements. To aid in data interpretation, hiPSC-CM MEA studies should report important technical details (e.g., pacing frequency, stimulation settings, cell culture duration) and should consider reporting raw data values to facilitate data comparison between studies.

Another critical parameter to consider when performing MEA experiments is the length of time hiPSC-CMs are grown in culture. hiPSC-CMs are immature cardiomyocytes with a fetal-like phenotype that are in a continued state of maturation (for review(14, 64)). As such, extended periods of cell culture have been shown to shift the phenotype of these cells to a more adult-like state (18, 56, 65). Between 1– vs 4-weeks of cell culture, Kumar et al. reported that older hiPSC-CMs have a more mature ultrastructure with increased myofibril organization, refined sarcomere structures, developed intercalated discs, and increased mitochondrial density (57). Older 4-week hiPSC-CMs also have increased expression for genes encoding cardiac ion channels (e.g., KCNJ2, KCNQ1, SCN5A) and calcium handling proteins (e.g., RYR2, CASQ2). In agreement, Nozaki et al. reported a stark shift in cardiac gene expression within the short time frame (1-15 days) recommended for MEA studies using a commercially available hiPSC-CM cell line (35). The latter included an 12-47-fold difference in key genes associated with voltage gated ion channel activity (e.g., KCNJ1, KCNQ1, CACNA2S2) and muscle contraction (e.g., CASQ2, MYBPC3). As such, passive maturation in cell culture is likely to influence hiPSC-CM electrophysiology under baseline conditions and in response to various drug compounds – which could also account for inter-facility variability in MEA studies. In the current two-week long study, we reported significant slowing of hiPSC-CM beating rate, increased depolarization spike amplitude, and lengthening of FPD and APD under standard cell culture conditions. These findings are in agreement with the study by Kumar, et al. which reported that extended cell culture time (4 weeks) slowed the beating rate and lengthened FPD in hiPSC-CMs, and Doss, et al. which reported longer APD90 in older hiPSC-CM (∼4 months)(66). In the current study, our analysis of contractile function was limited to excitation-contraction coupling delay time and contraction beat amplitude measurements. Even though these parameters were not changed during this two-week study, others have reported on the maturation of intracellular calcium handling within 15-30 days of culture (67) and larger calcium current density within 30-80 days of cell culture (56). Since we observed daily adaptations in hiPSC-CM physiology, it is beneficial to use these cells within a well-defined time range, and long-term cardiotoxicity studies may require additional planning.

Notably, we also observed that hiPSC-CM drug responsiveness is dynamic within the first two weeks of cell culture. As an example, younger hiPSC-CMs (4 days) were more responsive to E-4031 (I_Kr_ blocker) as compared to slightly older hiPSC-CMs (10 days of culture) – as demonstrated by a more significant lengthening of FPD and APD. In agreement, da Roche et al. reported that immature hiPSC-CMs with a “fetal-like” phenotype had an exaggerated response to E-4031 with longer APDs and an increased incidence of early delayed afterdepolarizations – as compared to “adult-like” hiPSC-CMs that underwent an extracellular matrix-mediated maturation protocol (68). This variable drug response may be attributed to the reliance of immature hiPSC-CMs on I_Kr_ for repolarization, as compared with older hiPSC-CMs (66). We also observed that older hiPSC-CMs (10 days) were more responsive to isoproterenol than younger hiPSC-CMs (4 days of culture), as demonstrated by a larger increase in the spontaneous beating rate and faster conduction velocity. This variable isoproterenol response could be related to maturation-dependent changes in both beta-adrenergic receptor expression and cAMP signaling (69, 70).

To improve preclinical cardiotoxicity testing, hiPSC-CM MEA experiments have been adopted as a vital component of the CiPA initiative. Implementing and rigorously reporting experimental techniques can improve reproducibility both within and between laboratories, and further support the utility of hiPSC-CMs as a high throughput screening tool. Toward this goal, our study highlights a few parameters to control for when planning hiPSC-CM MEA studies, including cell age, equilibrium duration, layout of drug treatments across a multiwell plate, and pacing stimulus settings. To date, the cardiac research field largely reports MEA measurements as a percent change from baseline, but also reporting raw data values will be help to improve data interpretation between published studies.

A few limitations in our study design should be considered and expanded upon by future work. First, the current study was limited to the use of a single hiPSC-CM cell line (#01434, Fujifilm Cellular Dynamics), which is a heterogeneous mixture of cardiomyocytes with ventricular, atrial, and nodal-like attributes (7, 71, 72). Other research groups have documented phenotypic differences between commercially available hiPSC-CM cell lines (e.g., Cor.4U hiPSC-CM have a feaster beating rate and shorter FPD compared to iCell) (27, 29, 31, 32). Accordingly, the results of our study may not directly translate to other cell lines. Second, our study was limited to the use of a single MEA device; therefore, our results may/not be replicated in alternative high-throughput screening systems (e.g., microelectrode array, impedance-based, optical-based). Third, we monitored hiPSC-CMs physiology throughout the first two weeks post-defrost, which aligns with the manufacturer instructions on the appropriate timeframe for experimentation. Nevertheless, hiPSC-CM maturation continues during prolonged cell culture, which can impact cardiac electrophysiology and contractility measurements at baseline and in response to drug or chemical exposure (56, 57, 73).

## Data Availability Statement

The authors confirm that the data supporting the findings of this study are available within the article and online supporting tables.

## IRB Statement

Not applicable.

## Author Contributions

DG and NGP designed experiments and drafted the manuscript text; DG and JP performed experiments; DG, JP, and NGP analyzed data, prepared figures, and approved the final manuscript.

