## Supplemental Tables for "hiPSC-CM Electrophysiology: Impact of Temporal Changes and Study Parameters on Experimental Reproducibility"

**Supplemental Table 1: Spontaneous human induced pluripotent stem cell-derived cardiomyocyte electrophysiology values across 20 minutes of equilibration in a microelectrode system.** BPM: beats per minute, FPD: field potential duration, stdev = standard deviation. Ordinary one-way ANOVA with multiple comparisons to minute 20 using Holm-Sidak correction. n =107-120 replicates per measurement group.

| Minute | Beat Rate (BPM) | | | FPD (ms) | | | Spike Amplitude (mV) | | | Conduction Velocity (mm/ms) | | |
|---|---|---|---|---|---|---|---|---|---|---|---|---|
| | mean ± stdev | p value | n value | mean ± stdev | p value | n value | mean ± stdev | p value | n value | mean ± stdev | p value | n value |
| 2 | 40.18 ± 4.75 | <0.0001 | 120 | 385.8 ± 19.8 | <0.0001 | 108 | 1.108 ± 0.298 | 0.7839 | 117 | 0.252 ± 0.085 | 0.7644 | 107 |
| 3 | 41.74 ± 3.40 | <0.0001 | 120 | 379.0 ± 19.6 | <0.0001 | 112 | 1.070 ± 0.297 | 0.9993 | 118 | 0.260 ± 0.097 | 0.9949 | 109 |
| 4 | 43.20 ± 4.06 | <0.0001 | 120 | 373.5 ± 21.0 | <0.0001 | 115 | 1.053 ± 0.295 | >0.9999 | 119 | 0.260 ± 0.092 | 0.9953 | 110 |
| 5 | 44.45 ± 4.47 | <0.0001 | 120 | 367.9 ± 21.3 | <0.0001 | 117 | 1.053 ± 0.291 | >0.9999 | 119 | 0.261 ± 0.092 | 0.9957 | 111 |
| 6 | 45.34 ± 4.75 | <0.0001 | 120 | 364.9 ± 21.0 | 0.0057 | 118 | 1.029 ± 0.288 | 0.9993 | 119 | 0.267 ± 0.094 | 0.9997 | 113 |
| 7 | 45.97 ± 4.91 | <0.0001 | 120 | 363.4 ± 20.7 | 0.0415 | 119 | 1.030 ± 0.285 | 0.9993 | 119 | 0.267 ± 0.087 | 0.9997 | 116 |
| 8 | 46.37 ± 5.02 | 0.0001 | 120 | 361.7 ± 19.0 | 0.2344 | 116 | 1.034 ± 0.296 | 0.9994 | 119 | 0.262 ± 0.084 | 0.999 | 114 |
| 9 | 46.72 ± 5.07 | 0.001 | 120 | 360.5 ± 19.5 | 0.5391 | 117 | 1.027 ± 0.297 | 0.9991 | 119 | 0.261 ± 0.085 | 0.999 | 113 |
| 10 | 46.94 ± 5.19 | 0.0036 | 120 | 360.8 ± 18.8 | 0.4463 | 119 | 1.034 ± 0.299 | 0.9994 | 119 | 0.258 ± 0.085 | 0.9883 | 111 |
| 11 | 47.12 ± 5.26 | 0.0087 | 120 | 361.7 ± 19.6 | 0.2227 | 119 | 1.038 ± 0.308 | 0.9996 | 119 | 0.258 ± 0.088 | 0.9869 | 111 |
| 12 | 47.37 ± 5.27 | 0.0278 | 120 | 360.9 ± 18.3 | 0.4029 | 119 | 1.050 ± 0.304 | >0.9999 | 119 | 0.259 ± 0.086 | 0.9907 | 113 |
| 13 | 47.55 ± 5.34 | 0.0551 | 120 | 359.9 ± 18.4 | 0.7163 | 119 | 1.042 ± 0.306 | 0.9997 | 119 | 0.264 ± 0.089 | 0.9994 | 114 |
| 14 | 47.82 ± 5.34 | 0.1452 | 120 | 359.2 ± 18.2 | 0.9053 | 119 | 1.046 ± 0.290 | 0.9999 | 118 | 0.263 ± 0.089 | 0.9992 | 115 |
| 15 | 48.01 ± 5.47 | 0.2349 | 120 | 359.1 ± 18.0 | 0.9311 | 119 | 1.050 ± 0.316 | >0.9999 | 119 | 0.259 ± 0.085 | 0.9897 | 113 |
| 16 | 48.29 ± 5.53 | 0.4309 | 120 | 358.8 ± 17.6 | 0.9649 | 118 | 1.047 ± 0.321 | >0.9999 | 118 | 0.258 ± 0.089 | 0.9892 | 110 |
| 17 | 48.46 ± 5.62 | 0.5099 | 120 | 358.6 ± 17.7 | 0.986 | 117 | 1.035 ± 0.300 | 0.9995 | 118 | 0.265 ± 0.094 | 0.9995 | 110 |
| 18 | 48.75 ± 5.74 | 0.6569 | 120 | 356.7 ± 17.9 | 0.9996 | 118 | 1.055 ± 0.333 | 0.9999 | 119 | 0.257 ± 0.087 | 0.9817 | 108 |
| 19 | 49.01 ± 5.82 | 0.6868 | 120 | 356.1 ± 17.8 | >0.9999 | 118 | 1.046 ± 0.316 | 0.9999 | 118 | 0.261 ± 0.090 | 0.9988 | 108 |
| 20 | 49.28 ± 5.91 | - | 120 | 356.1 ± 18.8 | - | 118 | 1.050 ± 0.319 | - | 118 | 0.269 ± 0.098 | - | 111 |

**Supplemental Table 2: Spontaneous human induced pluripotent stem cell-derived cardiomyocyte electrophysiology values across 14 days of culture.** BPM: beats per minute, FPD: field potential duration, stdev = standard deviation. Ordinary one-way ANOVA with multiple comparisons to day 7 using Holm-Sidak correction.

| Day | Beat Rate (BPM) | | | FPD (ms) | | | Spike Amplitude (mV) | | | Conduction Velocity (mm/ms) | | |
|---|---|---|---|---|---|---|---|---|---|---|---|---|
| | mean ± stdev | p value | n value | mean ± stdev | p value | n value | mean ± stdev | p value | n value | mean ± stdev | p value | n value |
| 2 | 53.80 ± 5.12 | <0.0001 | 148 | 288.9 ± 15.7 | <0.0001 | 141 | 0.301 ± 0.238 | <0.0001 | 167 | 0.328 ± 0.095 | <0.0001 | 63 |
| 3 | 49.91 ± 8.29 | <0.0001 | 264 | 352.6 ± 27.3 | <0.0001 | 248 | 0.860 ± 0.438 | <0.0001 | 257 | 0.291 ± 0.088 | <0.0001 | 224 |
| 4 | 48.32 ± 5.59 | 0.0001 | 363 | 372.1 ± 27.6 | <0.0001 | 351 | 1.162 ± 0.350 | <0.0001 | 358 | 0.262 ± 0.089 | 0.6867 | 344 |
| 5 | 49.48 ± 6.10 | <0.0001 | 138 | 412.8 ± 24.9 | <0.0001 | 134 | 1.701 ± 0.419 | <0.0001 | 135 | 0.207 ± 0.046 | <0.0001 | 133 |
| 6 | 46.30 ± 4.20 | 0.9112 | 228 | 435.1 ± 23.7 | <0.0001 | 223 | 2.023 ± 0.510 | <0.0001 | 232 | 0.252 ± 0.060 | 0.8214 | 203 |
| 7 | 46.22 ± 7.83 | - | 268 | 450.6 ± 39.5 | - | 255 | 2.452 ± 0.763 | - | 267 | 0.256 ± 0.060 | - | 240 |
| 8 | 45.89 ± 5.25 | 0.9112 | 68 | 465.2 ± 24.9 | 0.0012 | 62 | 2.707 ± 0.763 | 0.0026 | 68 | 0.258 ± 0.058 | 0.8528 | 62 |
| 9 | 42.98 ± 5.11 | 0.0005 | 66 | 499.7 ± 32.5 | <0.0001 | 61 | 2.637 ± 0.621 | 0.0192 | 67 | 0.286 ± 0.070 | 0.0265 | 63 |
| 10 | 39.41 ± 6.80 | <0.0001 | 75 | 505.9 ± 59.5 | <0.0001 | 82 | 2.691 ± 0.647 | 0.0026 | 86 | 0.353 ± 0.116 | <0.0001 | 74 |
| 11 | 39.04 ± 4.76 | <0.0001 | 68 | 547.9 ± 37.0 | <0.0001 | 65 | 2.987 ± 0.729 | <0.0001 | 68 | 0.329 ± 0.075 | <0.0001 | 61 |
| 12 | 39.64 ± 3.26 | <0.0001 | 34 | 566.4 ± 24.3 | <0.0001 | 33 | 3.303 ± 0.723 | <0.0001 | 34 | 0.298 ± 0.084 | 0.0236 | 32 |
| 13 | 40.07 ± 5.96 | <0.0001 | 68 | 550.0 ± 49.0 | <0.0001 | 64 | 2.838 ± 1.179 | <0.0001 | 72 | 0.347 ± 0.117 | <0.0001 | 60 |
| 14 | 41.12 ± 4.48 | <0.0001 | 40 | 545.9 ± 24.2 | <0.0001 | 36 | 3.611 ± 0.774 | <0.0001 | 39 | 0.296 ± 0.055 | 0.0236 | 37 |

**Supplemental Table 3: Paced (1.5 Hz) human induced pluripotent stem cell-derived cardiomyocyte electrophysiology values across 14 days of culture.** FPD: field potential duration, stdev = standard deviation. Ordinary one-way ANOVA with multiple comparisons to day 7 using Holm-Sidak correction.

| Day | FPD (ms) | | | Spike Amplitude (mV) | | | Conduction Velocity (mm/ms) | | |
|---|---|---|---|---|---|---|---|---|---|
| | mean ± stdev | p value | n value | mean ± stdev | p value | n value | mean ± stdev | p value | n value |
| 3 | 275.5 ± 13.8 | <0.0001 | 180 | 3.819 ± 1.109 | <0.0001 | 216 | 0.239 ± 0.024 | 0.195 | 204 |
| 4 | 282.1 ± 18.5 | <0.0001 | 300 | 4.150 ± 0.969 | <0.0001 | 313 | 0.225 ± 0.041 | 0.0144 | 318 |
| 5 | 311.5 ± 13.5 | <0.0001 | 132 | 4.269 ± 0.916 | <0.0001 | 124 | 0.204 ± 0.018 | <0.0001 | 134 |
| 6 | 316.6 ± 13.8 | 0.0268 | 164 | 5.012 ± 0.898 | 0.0023 | 142 | 0.220 ± 0.021 | 0.0006 | 137 |
| 7 | 320.9 ± 13.9 | - | 151 | 5.434 ± 1.254 | - | 145 | 0.233 ± 0.023 | - | 143 |
| 8 | 323.6 ± 14.6 | 0.2339 | 62 | 5.633 ± 1.092 | 0.1983 | 64 | 0.226 ± 0.029 | 0.2082 | 62 |
| 9 | 337.6 ± 14.3 | <0.0001 | 62 | 5.693 ± 0.829 | 0.176 | 65 | 0.230 ± 0.029 | 0.4402 | 63 |
| 10 | 333.7 ± 13.8 | <0.0001 | 55 | 5.940 ± 1.005 | 0.0038 | 62 | 0.256 ± 0.040 | <0.0001 | 60 |
| 11 | 332.6 ± 15.8 | <0.0001 | 43 | 6.596 ± 0.925 | <0.0001 | 41 | 0.254 ± 0.022 | 0.0002 | 42 |
| 12 | 344.4 ± 14.1 | <0.0001 | 32 | 6.548 ± 0.944 | <0.0001 | 34 | 0.265 ± 0.014 | <0.0001 | 33 |
| 13 | 333.9 ± 16.5 | <0.0001 | 46 | 6.690 ± 1.378 | <0.0001 | 48 | 0.280 ± 0.021 | <0.0001 | 40 |
| 14 | 336.0 ± 14.0 | <0.0001 | 38 | 6.871 ± 0.899 | <0.0001 | 38 | 0.275 ± 0.017 | <0.0001 | 35 |

**Supplemental Table 4: Paced (1.5 Hz) human induced pluripotent stem cell-derived cardiomyocyte action potential duration measurements from day 4 to 10 of culture.** APD30: action potential duration at 30% repolarization, APD50: action potential duration at 50% repolarization, APD90: action potential duration at 90% repolarization, stdev = standard deviation. Ordinary one-way ANOVA with multiple comparisons to day 7 using Holm-Sidak correction.

| Day | APD30 (ms) | | | APD50 (ms) | | | APD90 (ms) | | |
|---|---|---|---|---|---|---|---|---|---|
| | mean ± stdev | p value | n value | mean ± stdev | p value | n value | mean ± stdev | p value | n value |
| 4 | 171.7 ± 19.4 | 0.6239 | 15 | 247.6 ± 16.4 | 0.0213 | 15 | 319.6 ± 15.3 | <0.0001 | 15 |
| 5 | 169.5 ± 21.0 | 0.6239 | 10 | 257.3 ± 6.6 | 0.4237 | 9 | 336.7 ± 11.7 | 0.0004 | 10 |
| 6 | 169.5 ± 13.8 | 0.6239 | 9 | 253.0 ± 18.6 | 0.2404 | 10 | 352.1 ± 19.6 | 0.2333 | 10 |
| 7 | 181.5 ± 19.1 | - | 17 | 266.0 ± 15.8 | - | 17 | 359.2 ± 12.4 | - | 17 |
| 8 | 182.2 ± 20.6 | 0.9308 | 27 | 269.5 ± 16.5 | 0.522 | 27 | 365.9 ± 12.3 | 0.2333 | 27 |
| 9 | 184.5 ± 32.6 | 0.9308 | 10 | 279.1 ± 25.6 | 0.2404 | 10 | 389.7 ± 13.2 | <0.0001 | 10 |
| 10 | 203.7 ± 26.0 | 0.026 | 17 | 298.4 ± 20.5 | <0.0001 | 17 | 399.2 ± 15.5 | <0.0001 | 17 |
